# The use of variational autoencoders to characterise the heterogeneous subpopulations that arise due to antibiotic treatment

**DOI:** 10.1101/2024.12.19.629541

**Authors:** Dennis Bersenev, Emily Zhang

## Abstract

Antimicrobial resistance (AMR) is a persistent threat to global agriculture and healthcare systems. One of the challenges towards development of robust antimicrobials to date has been the limitation posed by low resolution bacterial sequencing technologies. The recent development of Bacterial Single Cell RNA sequencing protocols has provided an unprecedented opportunity in AMR research as it now enables researchers to probe bacterial populations at single cell resolution. In this study, we apply a Bayesian Variational Autoencoder, MrVI, to data generated by one such Bacterial Single Cell RNA sequencing protocol, BacDrop, and use it characterise changes in gene expression levels before and after antibiotic perturbation. Through the use of MrVI, we were able to find distinct DNA damage and heat shock response subpopulations. We also determined that each of the subpopulations could be mapped back to its respective antibiotic treatments, providing more precise insight into their mechanisms of resistance. These preliminary results indicate the potential that this new window into intracellular bacterial communication provides, and motivate the continued exploration of models to unveil the mechanisms underlying AMR.

## Introduction

Antimicrobial resistance is a pressing issue within the healthcare industry; a recent study published in the Lancet estimated that in 2019 alone, there were 4.95 million deaths associated with it.^19^ The driving causes of this rapidly emerging resistance include the inappropriate prescription of antibiotics and the extensive use of antibiotics as growth supplements in the agricultural industries worldwide.^20^ Moreover, the research into the development of new drugs has been significantly scaled back by major pharmaceutical companies in recent years, often being seen as a less profitable investment due to these drugs’ ^20^ short-term and often curative nature.

Ultimately, due to the pressing nature of this crisis, further research must be done in regard to the prevention of antibiotic resistance and the development of new drugs and treatment plans. In order to better characterise the heterogeneous stress response that bacteria experience after exposure to antibiotics, Ma et. al. introduced a novel sequencing protocol that can better detect subtle variations in gene expression at lower costs.^21^ A better understanding of the dynamic antibiotic-resistance responses that bacterial cells undergo allows for the development of highly targeted therapeutics with increased effectiveness.

The data from the BacDrop study was collected from an experiment which involved four different environmental conditions corresponding to four different sample sets, and two different replicates of all those sets^21^ . The standard approach to handling batch effects, specifically those involved in single-mode scRNA seq data where each batch contains common features, is through horizontal integration methods^10^ .

Horizontal integration methods aim at addressing the systemic deviations that arise when data are collected in batches, termed batch effects. The aim of batch effect correction is to remove the undesired technical variation introduced by differences in how sample data is collected across experiments while preserving true biological variation within and between experiments^10^ . The growing availability of reference single-cell atlases, such as the Human Cell Atlas^11^, has drawn significant attention to shortcomings in this step of the single-cell RNA sequencing (scRNA-seq) analysis workflow. Techniques such limma^12^ and ComBat^13^, have had disappointing results in the context of scRNA-seq data. These likely being due to their assumption of identical cell type composition across batches^1^ .

The issue concerning this assumption is that analyses involving scRNA-seq have been successful at demonstrating that even biological replicates, likely due to differences in sample collection and library preparation, have demonstrable differences in cell type composition^10^ .

In reconciling these shortcomings, newer, scRNA-seq adapted approaches to batch correction have at their core nonlinear batch-correction strategies^10^ . A prominent example, due to both its widespread popularity and its pertinence to our contrastive analysis with its utilisation in Bacdrop, is that of batch correction in Seurat v3 by means of mutual nearest neighbours (MNN)^1,10^ . MNN is an algorithm that can be used to find cells that are similar to one another across batches^1^, this is achieved by running k nearest neighbours (KNN) for each cell to find its nearest neighbours in all other batches, after repeating for all batches, cells which found each other as nearest neighbours become MNN pairs. The difference in expression values between cells in these MNN pairs can be averaged to provide an estimate of the batch effect. From this estimate, a correction vector can be obtained and used to map the original expression values to a common, batch-corrected, space (Figure 1)^1^ .

**Fig. 1.**
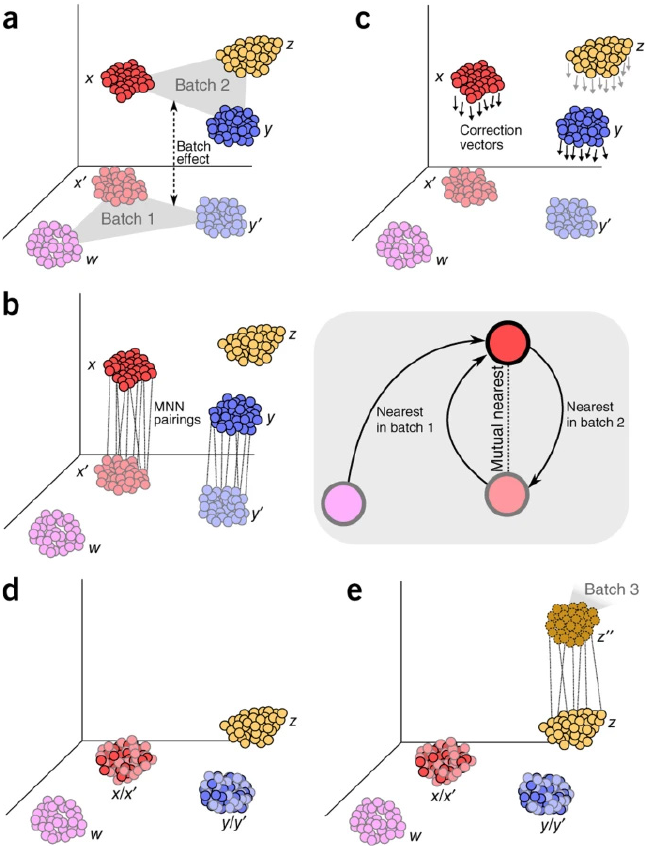
MNN Haghverdi, L., Lun, A., Morgan, M. et al. (2018)

The issue with common horizontal integration methods based on MNN, such as those used in the Seurat pipeline, lies in their continued inability to identify a shared representation able to account for the heterogeneity due to differences across both samples and individuals^14^ . In addressing these persisting challenges to solving the batch correction problem, a recent preprint from Boyeau et al.^14^ has proposed a multi-resolution deep generative model, MrVI, which is able to disentangle batch-related effects from differences in gene expression reflective of true biological differences.

Using MrVI with Bacdrop’s data as an exemplary case of multiple tiers of batch effects, we were able to learn a disentangled latent representation of the gene expression where all differences contained therein were reflective of true biological variability. We clustered the data obtained from the MrVI results and compared the various clusters to assign them to biologically significant subpopulations. These results were then compared to BacDrop’s Analysis. Ultimately, we were able to find an additional ‘heat shock response’ cluster and we were able to assign the clusters to specific antibiotic treatments.

## Methods

### Data

The single-cell RNA sequencing (scRNA-seq) data was procured from a study that introduces novel sequencing technology that aims to better characterise within-population bacterial heterogeneity. ^21^ The proposed scRNA-seq technology, BacDrop, has an rRNA-depletion step using RNase H and DNase I, “reducing sequence cost by at least 10-fold while simultaneously increasing information content”.^21^ This technique involves a cell fixation and permeabilization protocol before droplet encapsulation; this key step prevents cell lysis, thereby helping to isolate the mRNA of interest from the significantly more abundant rRNA.^21^ In the study, BacDrop was then used to characterise the heterogeneous subpopulations that arose after clinical isolates of *K. pneumoniae* were exposed to antibiotics.^21^ Ma et. al. analysed the data that was generated with the standard Seurat preprocessing pipeline, ultimately finding six distinct bacterial subpopulations.^21^ The aim of this project is to find a different but meaningful biological interpretation of this data using VAEs, quantifying sample-level heterogeneity within the bacterial subpopulations.

### Methodology

Variational Autoencoders (VAEs), are a type of unsupervised machine learning model concerned with the task of variational inference to estimate an intractable probability density function, this places VAEs under the category of explicit generative models^6^ . VAEs are based on a class of non-generative unsupervised machine learning models called autoencoders. Autoencoders are principally concerned with the task of dimensionality reduction: learning a low dimensional feature space to adequately represent an input data set^7^ . The autoencoder architecture consists of two components, encoder and decoder. The former mapping input data to a lower dimensional feature space, and the latter doing the inverse. The probabilistic spin on autoencoders, VAEs, coerce the decoder and encoder networks to estimate parameters of probability distributions, from which feature and data vectors can be sampled^6^; whereas, the autoencoder networks determine those vectors directly^7^ .

The application of VAEs typically comes in two varieties. Broadly, these are interpolation and compression. The interpolative character of VAEs manifests in their ability to generate data by drawing samples from a distribution approximating that which the input data is believed to follow. While compression comes in the form of the latent distribution. It is this second application of VAEs that has found the most success in the computational biology literature, given its ability to handle the large complex datasets one typically encounters^8,9^ .

The data we were interested in investigating, as outlined previously, was collected from an experiment which involved four different environmental conditions corresponding to four different sample sets, and two different replicates of all those sets^21^ . Naively concatenating the expression matrices from all these batches together was found to significantly distort downstream analysis (supplement 1). This result was unsurprising given the literature on the technical noise introduced by the process of sample collection itself^10^ . The standard approach to handling batch effects, specifically those involved in single-mode scRNA seq data where each batch contains common features, in this case those features being the same cell barcodes, is through horizontal integration methods^10^ . This was the approach taken by the authors of Bacdrop, and in an effort to enable a more robust analysis, using the powerful tool of VAEs, we used MrVI to learn a common metric space from raw expression counts to use as input for downstream analysis.

The MrVI model is introduced as a hierarchical probabilistic graphical model (Figure 2). The model is trained using the standard Bayesian Variational Autoencoder approach. In particular MrVI’s generative model, for each cell n, is as follows:

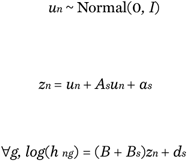

**Fig. 2.**
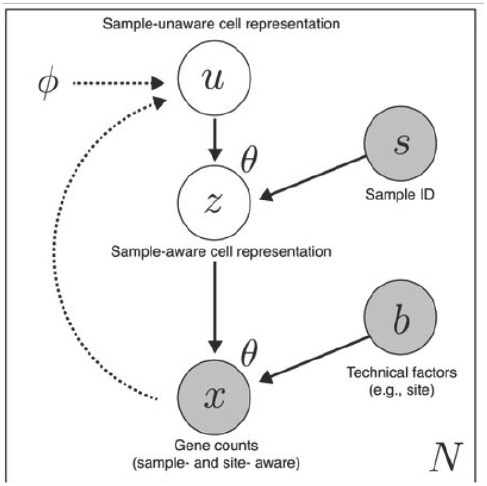
MrVI Probabilistic Graphical Representation

Where B captures site-agnostic expression patterns, *B*_*s*_ incorporates site specific expression patterns, and *d*_*s*_ is the column vector corresponding to sample s for cell n in its sample-sample distance matrix D(n), which has entries defined as:

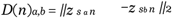

Lastly, normalised expression values are obtained:

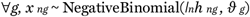

Where *I*_*n*_ is the size factor, defining the total sum of counts for cell n, and *ϑ*_*g*_ denotes the inverse dispersion parameter.

The latent space of interest, z, is obtained during inference according to a deterministic mapping, and therefore the VAE architecture has the structure seen in Figure 3. Wherein evidence lower bound (ELBO) is maximised to obtain a distribution approximating that of the intractable data likelihood, p(x).

**Fig. 3.**
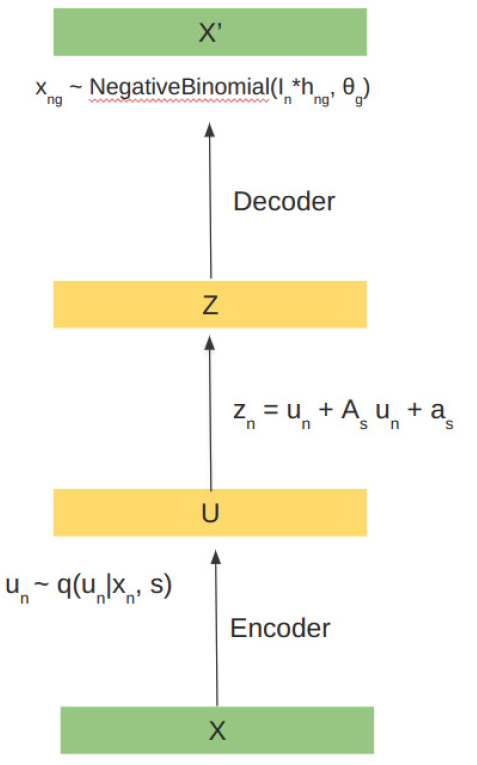
MrVAE

### Application

To obtain this latent space for the data provided in Bacdrop we first downloaded the MGH66 datasets, made available by the Bacdrop authors in the GEO repository: GSE180237. The raw gene count matrices representing these datasets were simply concatenated together by the union across genes (represented by their accession numbers), where each repeated gene was hyphenated to avoid the issue of duplicate variable names for downstream processing^5^ . The concatenation process incorporated meta information about sample and replicate of origin into the resultant data structure. After removing cells with low gene counts and genes with low cell counts, MrVI was trained using this data. Following training, the latent spaces were used for downstream analysis. To perform this analysis we made use of the tools available in the Scanpy package in Python^16^ . First, we further filtered cells expressing fewer than 40 genes, from which the neighbourhood graph was computed using both the u and z latent spaces. As expected, the visual provided by a 2D UMAP projection revealed how comparison in the z-space better accounted for sample variability (Supplementary Figure 2).

The latent feature representation was used to cluster cells and construct the neighbourhood graph using two different algorithms. The first based on k nearest neighbours (KNN), wherein cells were grouped according to their Euclidean distance from all other cells. The second grouped cells using a Gaussian kernel. With this approach, a cell’s neighbours are weighted using a Gausian kernel, with width given by k, and those cells more distant are assigned lower weights ^2^ . Using these neighbourhood graphs, the Leiden algorithm ^3^ was then applied to partition the different communities. Comparison between these approaches is in (Supplementary Figure 3).

## Results

### Seurat Comparison

Different clustering methods were explored in the analysis step to determine a meaningful representation of the within-population heterogeneity that arose with exposure to ciprofloxacin, gentamicin, and meropenem.

This naive clustering approach identified 53 distinct subpopulations that were clustered into two larger subpopulations (Supplemental figure 1b). The untreated cells are largely clustered together and fall into many of the same subpopulations but the treated cells are spread out and interspersed between the various subpopulations (Supplemental figure 1a). This could indicate that the *K. pneumoniae* isolates responded similarly to the three different treatments. However, the relatively large number of clusters that were produced with this approach greatly overwhelms the six that were identified in the BacDrop study^21^ and the separation of the clusters themselves is quite poor and difficult to distinguish from one another (Supplemental figure 1b). In order to produce results that allow a more meaningful biological interpretation of each of the subpopulations, we used more sophisticated approaches.

**Figure 1:**
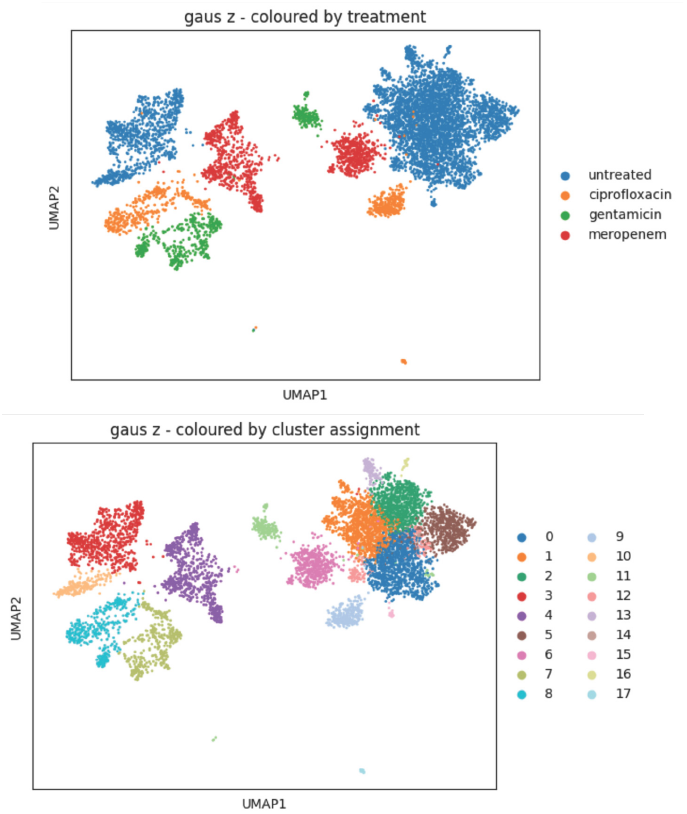
The z latent space with Gauss Clustering Approach a) UMAP plot for the z latent space with Gauss clustering, coloured by treatment grouping b) UMAP plot for the z latent space with Gauss clustering, coloured by cluster assignment (arbitrarily labelled)

In general, clustering data in the z latent space produced more distinct and clearly separated clusters regardless of the approach (Supplemental figure 2b, Figure 1b). As such, we decided to move forward with the z latent space for our downstream analysis. Moreover, within the z latent space, the Gaussian clustering approach produced a more manageable and realistic number of clusters; with nearly half the number of clusters of the KNN approach (Figure 1b, Supplemental figures 3b), finding the biological significance of each of these clusters and assigning them to heterogeneous subpopulations will be considerably more straightforward.

The data was then logarithmized and the relative gene expression levels of each cluster were ranked with Scanpy’s ‘rank gene groups’ function (Supplemental figure 3).^22^ The top 25 highly expressed genes for the clusters that were exposed to antibiotics were analysed to determine their biological function and marker genes for each of the subpopulations were determined (Supplemental table 1). This was done by querying the NCBI Protein database and the Interpro database with the accession number of each of these genes, and conducting literature reviews for genes of interest to determine if they can confidently be classified as biological markers. The bacterial subpopulations that were subsequently identified include MGE (mobile genetic elements), DNA replication, heat shock response, DNA damage response, multidrug efflux, and protein damage response (Figure 2). Clusters 3, 4, 7, 8, and 10 were identified to be a part of the MGE cluster, with the majority of cells upregulating 3 different transposase proteins (Figure 2).

Transposons play a significant role in bacterial antibiotic resistance as they are able to convert transcriptionally silenced dormant genes into determinants of antibiotic resistance by introducing promoters into random genomic locations.^23^ However, the clusters that were associated with MGE were difficult to distinguish from each other, having very similar expression patterns of very similar proteins.

**Figure 2:**
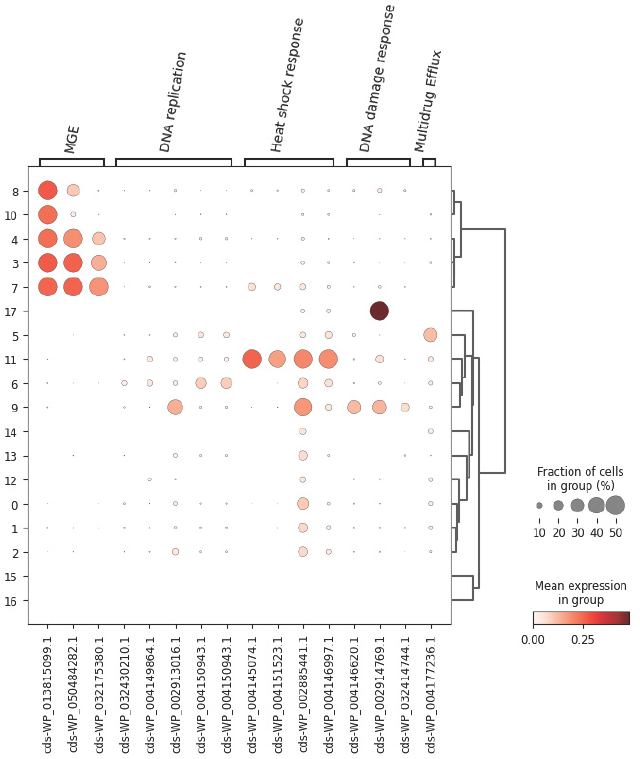
The gene expression values corresponding to the Gauss clusters in the z latent space, ordered by their respective bacterial subpopulations

Moreover, cluster 17 has higher expression levels of RecA, which is a recombinase protein that is associated with the DNA damage response (Figure 2). This protein facilitates the repair of double-stranded breaks in DNA, coating single-stranded ^26^ DNA to form RecA filaments in the repair pathway.

The majority of cells in cluster 11 expressed the marker genes associated with a heat shock response, upregulating GroEL, DnaK, IbpA, and IbpB (Figure 2). GroEL and DnaK are chaperone proteins that assist with the folding of newly synthesised proteins during heat shock and with the repair of proteins that have been damaged by heat shock.^24^ These proteins may also act as innate immunostimulators, activating other stress-related proteins that could provide protection to the bacterial cells. Similarly, IbpA/IbpB are also involved in the bacterial heat shock response.^25^ This specific subpopulation was not isolated in the BacDrop study, but it is clearly distinguished from the other clusters in terms of gene expression levels with our approach.

Cluster 9 consisted of cells with the gene expression of GroEL, UvrA, and RecA upregulated. This likely corresponds with the DNA damage response. UvrA binds with UvrB to form a complex that is able to sense DNA damage in the cell, thereby allowing other proteins to repair the genome.^27^ While GroEL is a heat shock protein, it can also be upregulated in response ^24^ to general stress within the cell.

We were unable to characterise cluster 6 due to a lack of clearly identifiable markers. The genes that were highly expressed mainly consisted of genes encoding proteins for transcription, translation, and general bacterial metabolism. The marker genes were only expressed in a small fraction of the cells and the mean expression for each of these genes is also very low. As such, we decided to label this cluster as ambiguous.

The rest of the clusters were associated with the untreated group. Because we are examining the response of bacteria to antibiotic treatment, we did not examine the other clusters in depth.

**Figure 3:**
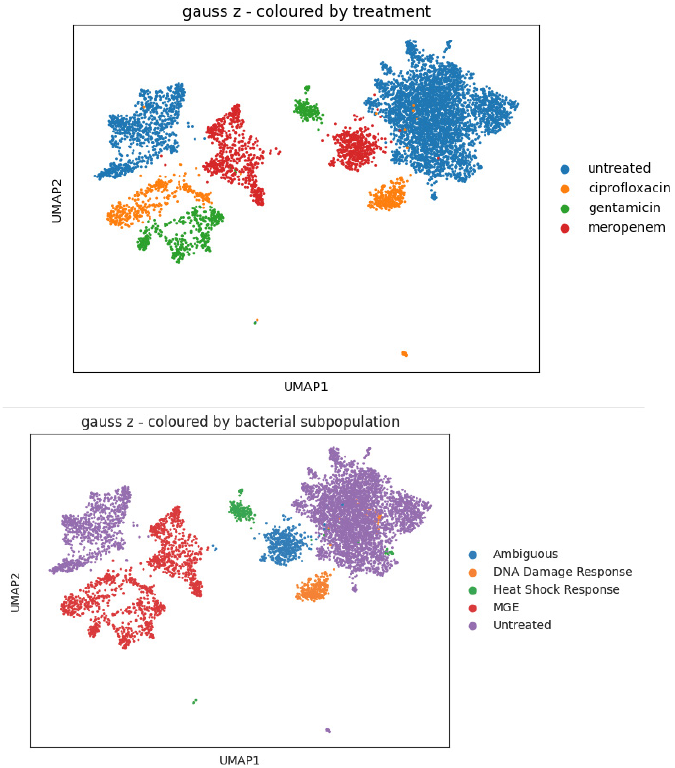
The bacterial cells with the z latent space, with Gauss clustering, coloured by their determined bacterial subpopulation and treatment a) UMAP plot for the z latent space with Gauss clustering, coloured by subpopulation b) UMAP plot for the z latent space with Gauss clustering, coloured by treatment (same as figure 1a for better visualisation)

After assigning the bacterial cells to their respective subpopulation, we can then assign the subpopulations to the treatment groups. The bacteria treated with gentamicin upregulated the expression of genes associated with the heat shock response (Figure 3a, 3b). On the other hand, the populations treated with ciprofloxacin showed higher expression of genes associated with the bacterial DNA damage response (Figure 3a, 3b). As well, regardless of treatment, all treatment options were associated with the MGE clusters, which corresponds to the upregulation of transposase enzymes.

## Discussion

### Conclusion

Upon applying a novel deep-generative model called MrVI to scRNA-seq data that was obtained from clinical isolates of *K. pneumoniae* before and after exposure to antibiotics, we were able to find several heterogeneous subpopulations that arose after treatment. While most of these subpopulations were identified in the BacDrop paper, we were able to distinguish between different clusters that correspond to DNA damage response and heat shock response. As well, we were able to associate the bacterial subpopulations with specific antibiotic treatments. This distinction between the bacterial response to the three different antibiotics could better elucidate the exact mechanisms that cells use to respond to perturbation. Further insight into these mechanisms that result in changes to gene expression levels could allow researchers and pharmaceutical companies to develop more highly targeted therapeutics with increased efficiency and decreased side effects. However, we were ultimately unable to detect some of the distinct subpopulations that were identified in the BacDrop paper. Perhaps with better optimization of the algorithms or more exploration of the different clustering techniques, we would be able to better locate and label the distinct bacterial subpopulations.

### Extensions

Something unaddressed in the model used for the present analysis concerns hyper-parameter selection. For VAEs this is the dimensionality of the latent feature space^6^, which was selected as MrVI’s default of 10^14^ . This decision was somewhat arbitrary, based on domain expertise of the hypothesised factors of variation. This selection motivated by the challenge of accurate selection of this hyperparameter presents^18^, in addition to the scalability issues involved in the compute heavy training cycle.

An alternative formulation of MrVI’s architecture could incorporate the Beta VAE approach proposed by Higgins et al^17^ . This approach provides a grounded methodology to learning disentangled latent representations through the use of *β* a hyperparameter which emphasises learning statistically independent latent factors^17^ . Finally, a recent method, multiGroupVI, tackles the problem of disentanglement by training encoders for each different experimental group individually, as well as a shared encoder to account for both the variation spanning groups and that which is group-specific^15^ . Perhaps such an approach could have greater success in determining the drivers of antibiotic resistance, than the methodology presented in this work.

## Acknowledgments

We would like to thank the authors of MrVI for their software’s open distribution, Pierre Boyeau for his correspondence concerning MrVI’s VAE architecture, the authors of Bacdrop for making their data accessible to the public, the developers of the scverse ecosystem for making high quality tools freely available, Dr. Jiarui Ding for his suggestions throughout the development of this work, and our colleagues and classmates for their attention and feedback during a premature demonstration of the present work.

## Respository

Github Repository: https://github.com/Dennis-Bersenev/GeneratingResistance

## Supplementary Figures

**Supplemental Figure 1:**
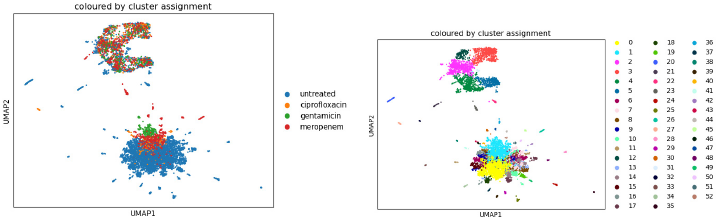
Naive normalisation grouping a) UMAP plot with naive normalisation resultant grouping, coloured by treatment grouping b) UMAP plot with naive normalisation resultant grouping, coloured by cluster assignment (arbitrarily labelled from 0 to 52)

**Supplemental Table 1:**
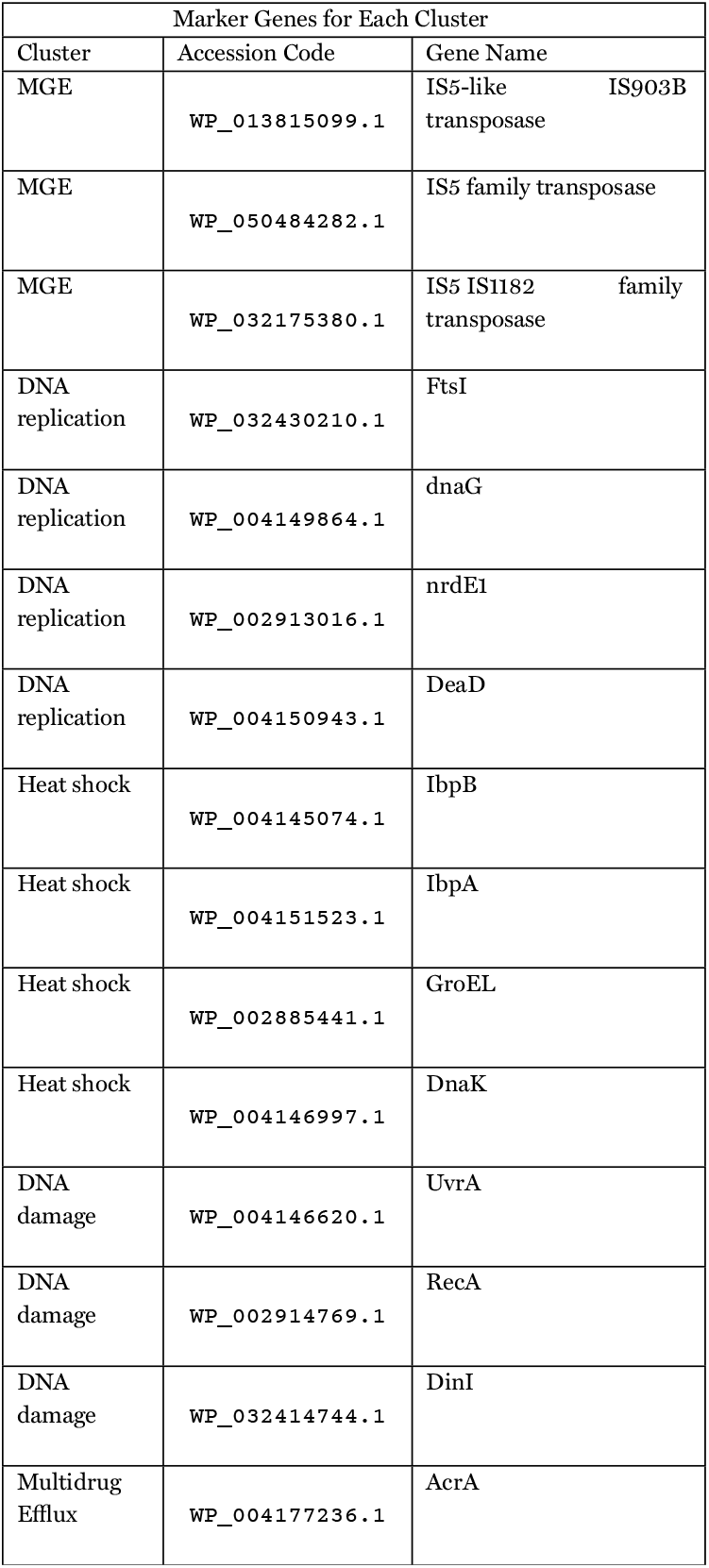
Marker genes for each bacterial subpopulation.

**Supplemental Figure 2:**
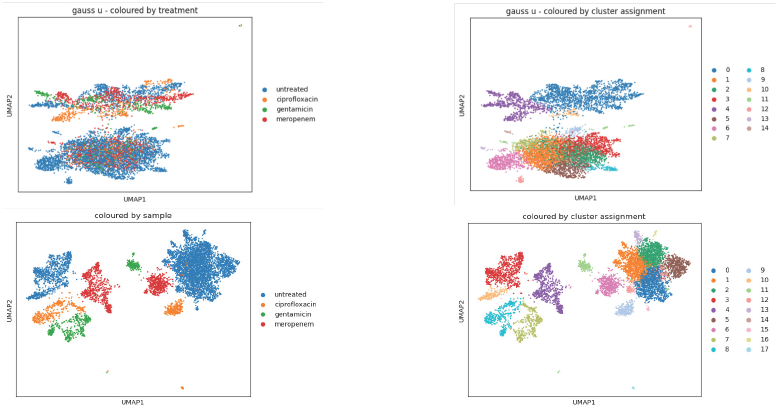
Clustering of sample-unaware latent space. a) UMAP plot for the u latent space with Gauss clustering, coloured by treatment grouping b) UMAP plot for the u latent space with Gauss clustering, coloured by cluster assignment (arbitrarily labelled) c) UMAP plot for the z latent space with Gauss clustering, coloured by treatment grouping d) UMAP plot for the z latent space with Gauss clustering, coloured by cluster assignment (arbitrarily labelled)

**Supplemental Figure 3:**
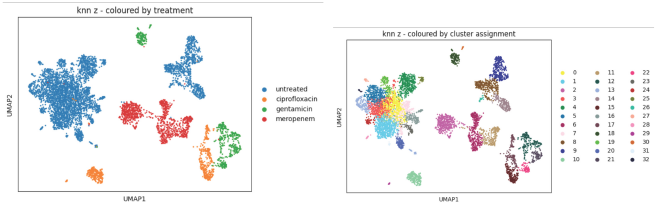
The z latent space with KNN Clustering Approach a) UMAP plot for the z latent space with KNN clustering, coloured by treatment grouping b) UMAP plot for the z latent space with KNN clustering, coloured by cluster assignment (arbitrarily labelled)

**Supplemental Figure 4:**
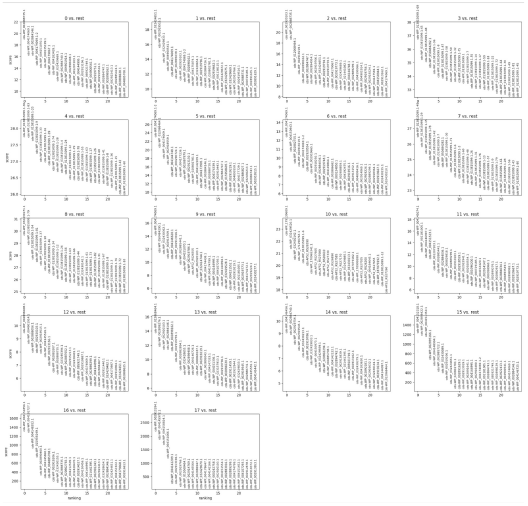
Top 25 highly expressed genes for each Gauss cluster in u latent space

